# PRISM: A Python Package for Interactive and Integrated Analysis of Multiplexed Tissue Microarrays

**DOI:** 10.1101/2024.12.23.630034

**Authors:** Rafael Tubelleza, Aaron Kilgallon, Chin Wee Tan, James Monkman, John Fraser, Arutha Kulasinghe

**Affiliations:** Frazer Institute, Faculty of Medicine, The University of Queensland, Brisbane, QLD, Australia; Queensland Spatial Biology Centre, Wesley Research Institute, The Wesley Hospital, Brisbane, QLD, Australia; Division of Bioinformatics, Walter and Eliza Hall Institute of Medical Research, Melbourne, VIC, Australia; Critical Care Research Group, The Prince Charles Hospital, Brisbane, QLD, Australia

## Abstract

Tissue microarrays (TMAs) enable researchers to analyse hundreds of tissue samples simultaneously by embedding multiple samples into single arrays, enabling conservation of valuable tissue samples and experimental reagents. Moreover, profiling TMAs allows efficient screening of tissue samples for translational and clinical applications. Multiplexed imaging technologies allow for spatial profiling of proteins at single cell resolution, providing insights into tumour microenvironments (TMEs) and disease mechanisms. High-plex spatial single cell protein profiling is a powerful tool for biomarker discovery and translational cancer research, however, there remain limited options for end-to-end computational analysis of this type of data. Here, we introduce PRISM, a Python package for interactive, end-to-end analyses of TMAs with a focus on translational and clinical research using multiplexed proteomic data from the CODEX, Phenocycler Fusion (Akoya Biosciences), Comet (Lunaphore), MACSima (Miltenyi Biotec), CosMx and Cellscape (Bruker Spatial Biology) platforms. PRISM leverages the SpatialData framework to standardise data storage and ensure interoperability with single cell and spatial analysis tools. It consists of two main components: TMA Image Analysis for marker-based tissue masking, TMA dearraying, cell segmentation, and single cell feature extraction; and AnnData Analysis for quality control, clustering, iterative cell-type annotation, and spatial analysis. Integrated as a plugin within napari, PRISM provides an intuitive and purely interactive graphical interface for real-time and human-in-the-loop analyses. PRISM supports efficient multi-resolution image processing and accelerates bioinformatics workflows using efficient scalable data structures, parallelisation and GPU acceleration. By combining modular flexibility, computational efficiency, and a completely interactive interface, PRISM simplifies the translation of raw multiplexed images to actionable clinical insights, empowering researchers to explore and interact effectively with spatial omics data.

## Introduction

Spatial proteomics, recently named the 2024 “Method of the Year” [1], provides deep insights into tissue architecture, enabling the construction of cellular and organisational atlases of normal tissues and disease biology. This technique is particularly relevant for our understanding of tumour progression. Spatial proteomics covers a wide range of methods including cyclic immunofluorescence (CycIF), co-detection by indexing (CODEX), and imaging mass cytometry (IMC). These approaches produce high-dimensional multiplexed imaging data of tissues, which can rapidly become a challenge to computationally analyse without dedicated interactive tools, especially for biologically focused scientists [2].

Tissue microarrays (TMAs) are a common tool for clinical research and enable researchers to profile patient tissues in a relatively cost-effective, yet high-throughput manner. Moreover, TMAs enable the screening of biomarkers across clinical cohorts, reducing slide-to-slide variability and reducing batch effects [3]. With the advent of multiplexed imaging technologies, such as CycIF and Phenocycler-Fusion (PCF or CODEX) (Akoya Biosciences, USA), it is possible to detect the in-situ spatial localisation of proteins from single cell to subcellular resolutions. Such capabilities have provided deeper insights into the features and mechanisms driving clinical endpoints such as progression free survival (PFS) and overall survival (OS) [4] [5] [6]. However, the limiting bottleneck for translating these insights to the clinic are the computational challenges in processing and analysing these high-dimensional imaging-based datasets [7].

Drawing meaningful biological insights from multiplexed images requires a broad processing and analysis pipeline encompassing different data modalities and types, requiring systematic methodologies. Starting from the raw multiplexed image, visual inspection is used to perform quality control (QC) on each marker for appropriate staining patterns and intensities.

Uniquely for TMAs, this is followed by de-arraying, which is the process of identifying the grid-like arrangement of tissue cores on the slide. This is followed by assigning each individual core a grid label, which is used to map back to the patient identifiers. Following this, cell segmentation is performed on the image. Current strategies generate cell segmentation masks by either 1) segmenting nuclei using a nuclear stain such as DAPI followed by an expansion of the nuclear mask to estimate cell boundaries, or 2) segmenting the entire cell directly using cytoplasmic markers like E-cadherin. Each strategy comes with its advantages and drawbacks. The former produces consistent results as nuclear markers usually have more ubiquitous and robust staining across tissues, with the drawback of not capturing the true morphology and bounds of the cell. The latter has the opposite, where a more realistic representation of the cell is captured, at the expense of consistent results due to the lack of ubiquitous and truly cytoplasmic markers across all cell types. Human-in-the-loop interactivity by visual inspection is therefore necessary to guide the choice of segmentation strategy. With the cell segmentation masks, image properties such as the mean or median marker intensities and morphologies for each cell are measured, effectively creating a cell-level feature table. Methods from the single cell bioinformatics space can then be applied to this feature table to perform marker normalisation, filtering, QC and cell type annotation.

Normalisation of intensity-based single cell feature tables remain a topic of debate, although inverse arcsine and log transforms have been recommended [8]. Uniquely, filtering and QC can be done across all modalities utilising the raw image, the de-arrayed cores, the cell segmentation masks, and the single cell feature table. After filtering and QC, cell types can then be annotated with 1) a user-guided approach involving unsupervised clustering followed by manual user annotation, or 2) an unguided approach involving machine learning-based transfer methods (STELLAR, CELESTA, etc. [9] [10]) which automatically assigns cell type labels. Next, spatial analyses are performed by utilising both the spatial coordinates and the cell type annotations of each cell to capture spatially informed neighbourhoods and interactions. Ultimately, these features can be used to compare against different clinical endpoints or groupings (e.g. response to therapy or overall survival data) to answer specific research questions [11] [12].

Existing software, tools, and pipelines for multiplexed images are limited in scope, require specific data formats and structures, lack user interactivity, scalable and fast visualisation, and are fragmented across different programming languages. The QuPath [13] software provides an entry point for multiplexed TMAs, providing visualisation capabilities, an interactive interface, a de-arrayer, but lacks comprehensive single cell level analyses and requires laborious data exports into other packages with such capabilities. MCMICRO [14] provides a pipeline covering most processing steps from raw image to spatial analysis, but lacks interactivity and user supervision to optimise parameters at each intermediate step. Furthermore, MCMICRO requires images and tables stored in specific formats and folder structures, reducing ease-of-access and interoperability. On the other hand, the Sopa pipeline [15] addresses this by adopting the newer SpatialData framework to store all data modalities in a standardised and interoperable format, but it also suffers from user interactivity at each intermediate step. SPArrOW [16] builds upon these shortcomings and offers an interactive and human in the loop solution but is tailored for imaging-based transcriptomic datasets. To our knowledge, there are no solutions that provide user interactivity across the entire scope of analysing raw, high-dimensional spatial proteomic data that is well-situated to provide fast clinical insights.

Here, we present PRISM, a Python package for Interactive and Integrated Analysis of Multiplexed Tissue Microarrays. With PRISM, we provide an accessible, scalable, and end-to-end solution that parses raw multiplexed TMAs while being well situated to correlate spatial single cell features to clinical outcomes, all in a single interactive graphical user interface (GUI).

## Main

### PRISM provides a highly user-interactive solution for processing and analysing multiplexed tissue microarrays

The motivation for PRISM was to make multiplexed TMA processing and analysis more interactive, accessible, and convenient for users. To this end, PRISM adopts and integrates specific data objects, tools, and graphical user interface frameworks. PRISM adopts SpatialData [17] as the core data object, using it as a container to incrementally add and store intermediate data derived either directly from the raw image, or from other intermediate data object(s). This allows for transparent and reproducible analyses with an analysis start-point possible at any point in the analysis pipeline. SpatialData also standardises the storage of these data formats, making data produced with PRISM accessible and interoperable with existing community led tools for single cell [18] [19] and spatial analysis [20] [21]. PRISM organises and implements these tools within modular computational components, each of which has a dedicated widget that the user interacts with inside the napari viewer [22] (Figure 1a). For visualisation, PRISM integrates with napari-spatialdata, the dedicated napari plugin for SpatialData objects.

**Figure 1:**
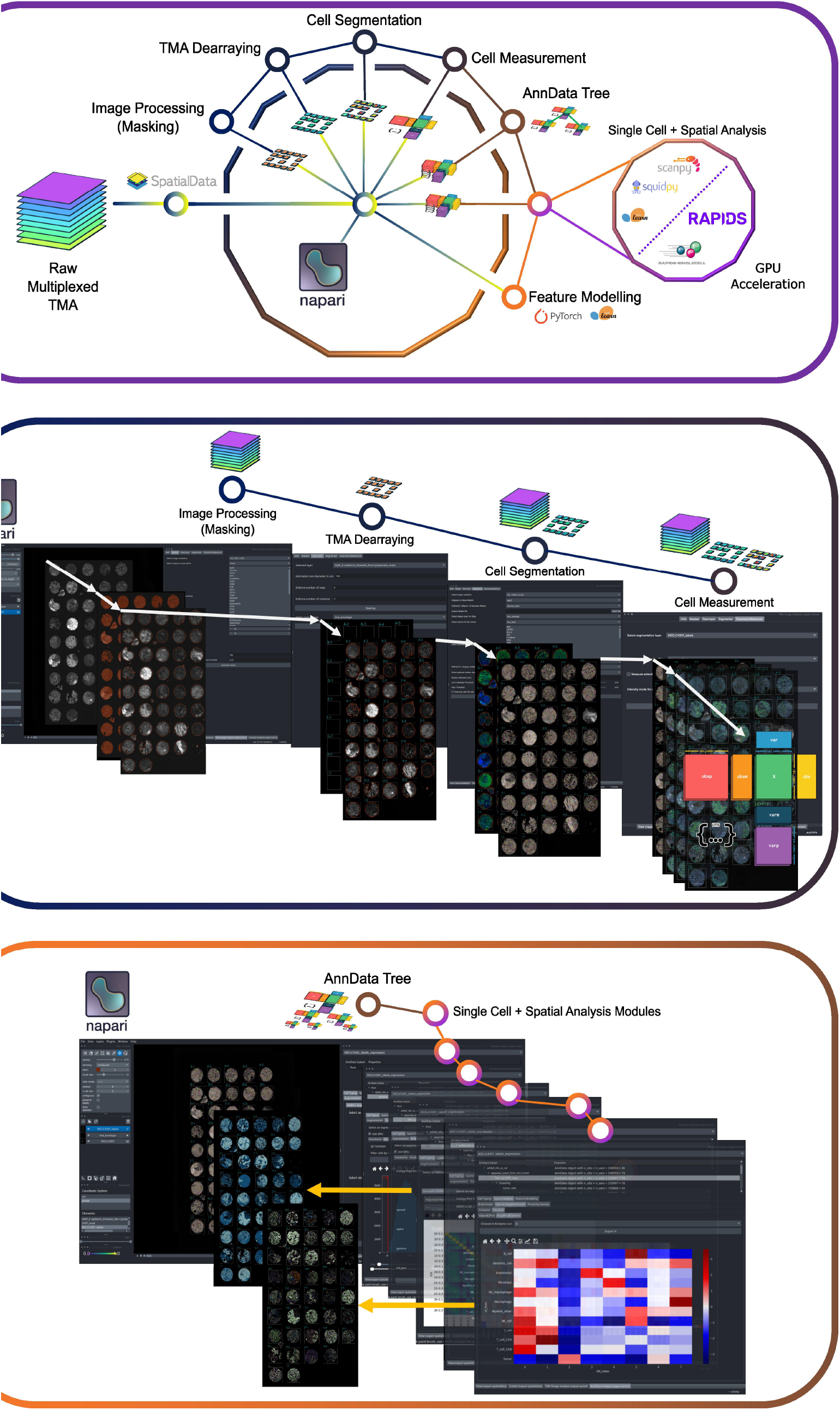
High level overview of PRISM, a multifaceted solution for interactive processing and analysis of multiplexed microarrays. **a)** Computational components of PRISM are structured and organised in a modular yet integrated fashion, adopting the SpatialData format to integrate with existing single cell and spatial analysis tools like Scanpy and Squidpy (GPU accelerated with rapids-singlecell and RAPIDS), as well as providing user-interactive interfaces within the napari viewer. **b**) The TMA image analysis group contains modules for processing and analysing raw multiplexed images into single cell expression AnnData tables. **c)** The AnnData analysis group facilitates interactive analyses by organising all tables and subsets into an interactive hierarchical tree that the user can select any AnnData node to analyse. The selected node can then be analysed by any of the provided single cell and spatial analysis modules. Analysis examples shown in this figure are multiplexed proteomic data from CODEX (PhenoCycler-Fusion) lung tissue samples.

PRISM contains two groups of components that ‘operate’ on specific data modalities contained within the SpatialData object (Figure 1a). The TMA image analysis group (Figure 1b) consists of four main modules. In the first module, the user chooses and tweaks a set of image processing functions to generate binary masks from a user-chosen set of marker channel(s). These masks can be used in the second module that provides an interface for dearraying or segmenting the TMA into distinct cores. The third module provides an extensive user interface for performing and optimising cell segmentation with Cellpose [23], with additional tools to generate pseudo-images by merging multiple markers for segmentation input and previewing input images. This module also provides utility for the implementation of custom-trained segmentation models. The last module uses cell segmentation masks to measure the marker channel intensities and morphological characteristics of every cell, then stores these into a single cell AnnData feature table [24]. The user can then directly analyse these in the same viewer with the AnnData analysis group. The AnnData analysis group contains an intermediate data module that stores all AnnData objects in a hierarchical tree structure, and a set of single cell and spatial analysis modules that operate on those objects. The analysis modules cover both conventional single cell bioinformatics to spatial analysis methods, where the user can perform feature table normalisation, filtering, clustering, cell type annotation, spatial graph generation, CN identification [25], and score the spatial proximity of cells [21]. Crucially, at every analysis step, the user can perform robust quality control and exploration by visualising cellular features and labels on the linked cell segmentation mask, alongside the raw image and de-arrayed TMA cores (Figure 1c). For performance and scalability, data are parsed as chunked Dask [26] arrays and on-disk Zarr [27] stores, allowing for lazy, asynchronous, parallelisable and GPU-accelerable computations and rendering. In particular, PRISM stores the raw TMA in a pyramidal image format, allowing for efficient visual rendering and image processing.

### PRISM Provides an Interface for Customisable and Interactive Image Masking, Identification and Annotation of TMA Cores, and Cell Segmentation

In the TMA image analysis group, PRISM provides a set of tools necessary for converting raw multiplexed TMA images into single cell level feature tables. This group consists of four main modules which cover masking, TMA dearraying, cell segmentation, and cell measurement. In the masking module, a user can generate masks representing the entire core or even regions within the core. PRISM gives the option for users to perform masking on a single or multiple channels, and at various resolutions (Figure 2a, middle panel). This gives the user control, tailoring masks to their desired application. For example, lower resolution masks that compute faster but more pixelated are suitable for coarser operations that operate on macroscopic objects in the image, such as TMA cores (Figure 2a, right panel). On the contrary, higher resolution masks that are computed slower but have more detail boundaries are suitable for tissue compartment annotations which may be used for further analysis (Figure 2a, right panel). Once the input image(s) are chosen, they can be processed using a variety of thresholding, correction, and noise-reduction filters, including Li thresholding, 2D Sobel filtering, contrast limited adaptive histogram equalisation (CLAHE), and gamma correction. Gaussian blurring is then applied to remove high wavelength spatial noise before Otsu intensity thresholding binarises the image to produce masks of the whole core and of each channel (Figure 2a, middle panel).

**Figure 2:**
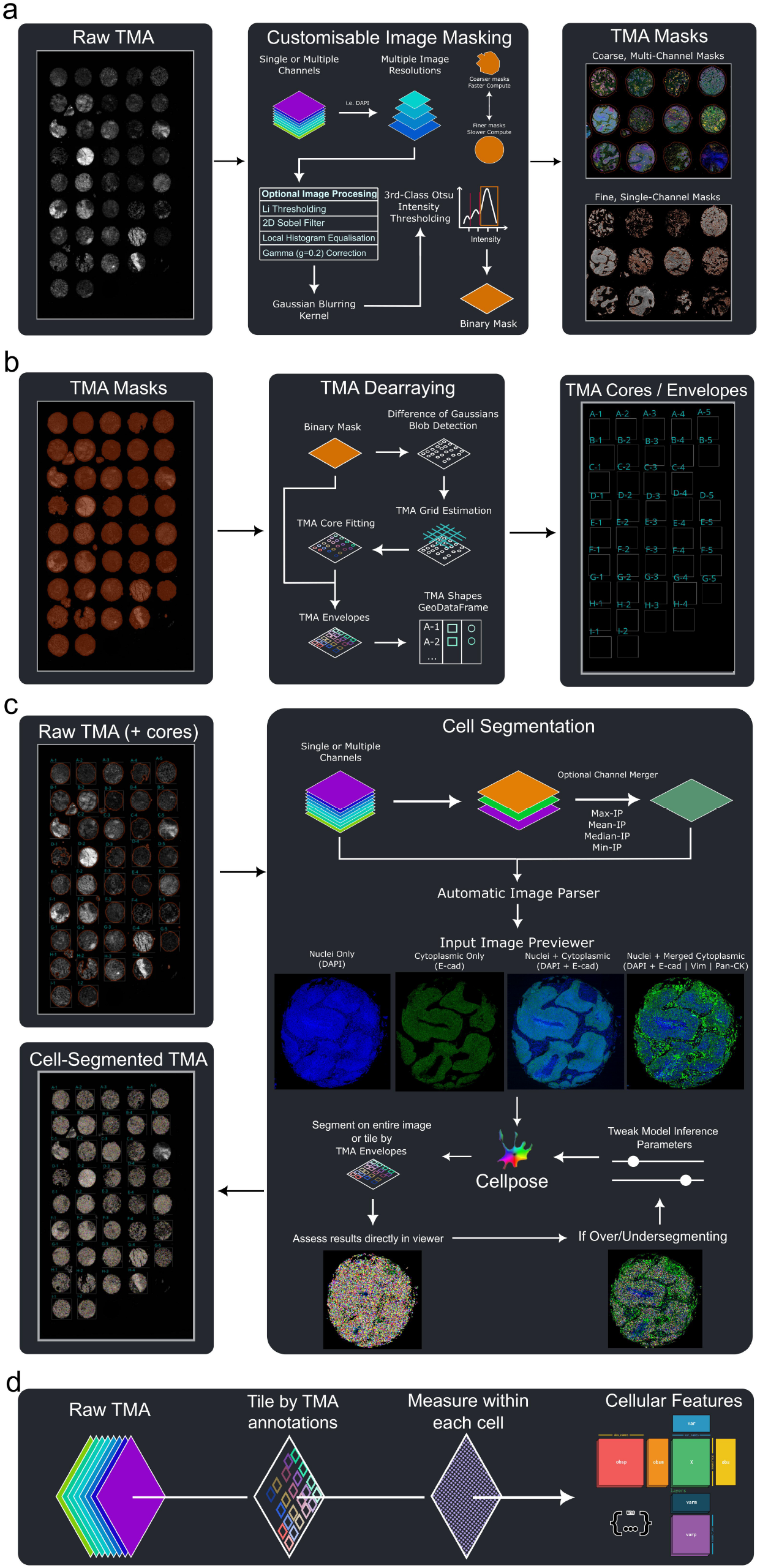
End-to-end demonstration of the TMA Image Analysis modules showing functionality for preprocessing, masking, dearraying, and segmentation on a multiplex-imaged TMA of Non-Small Cell Lung Carcinoma patient samples. **a)** The TMA masking algorithm is shown here: a raw multiplexed image from tissues samples imaged by the PhenoCycler-Fusion can be preprocessed (left panel) with a variety of noise-reduction methods (middle panel) before being thresholded into a set of whole-core masks (right, top panel) and single-channel masks (right, bottom panel). **b)** The binarised masks (left panel) are then de-arrayed by blob fitting and estimated grid positions (middle panel) are annotated onto the core label (right panel). **c)** The cell segmentation module ingests the raw TMA, and core masks, and it provides utility for channel merging to ensure optimal segmentation performance in the user’s chosen Cellpose segmentation algorithm (right panel, top row). The ingested channels are parsed for a given set of choice markers, evaluated by the chosen segmentation algorithm, and results are visualised for assessment and further tweaking by the user (right panel, bottom row).**d)** The last module performs image measurements on the segmentation masks tiled optionally by TMA annotations and outputs a single cell level AnnData table with the marker intensities, morphological measurements, and spatial coordinates of each cell.

TMA dearraying is typically a highly manual process that involves the manual region-of-interest selection and tissue truncation. In the de-arraying module, PRISM provides a fast method to de-array and annotate tissue cores via a core fitting routine that searches for blobs in the binary masks generated in the previous module, estimates an expected grid structure, and fits the blobs to this grid. PRISM stores cores annotations in two representations: 1) the conventional circular TMA core shapes, and 2) TMA envelopes. TMA envelopes loosen the constraint that cores are perfectly circular to capture irregularly shaped cores. This was done to capture edge cases of core geometry, which may be highly irregular in shape and/or partially fragmented. Users can then perform interactive quality control to resize, move, and delete TMA annotations in the napari viewer.

In the cell segmentation module, PRISM provides an interface for highly customisable Cellpose-based cell segmentation. Depending on the user’s dataset, they can choose to perform segmentation using a single channel, or two image channels for nuclei and cytoplasm. PRISM provides the functionality for easily merging multiple channels with maximum, mean, median and minimum intensity projections to jointly represent regions-of-interest (e.g. cytoplasm). All choices by the user at this point are automatically parsed into image formats and shapes that are directly compatible with Cellpose. Before segmentation, the user can assess their processed data by directly visualising the input images. The users can tile by TMA annotations made in the previous module, segmenting only within those annotations, and thereby skipping over background. The user can then directly assess and validate the output segmentation masks, tweak parameters and, re-run segmentation to further optimise the results, if desired. To finally bridge imaging data into single cell data, a simple interface is provided where the user can measure image properties of every cell in the segmentation mask. For each cell, the mean or median intensity of each marker channel can be measured as a proxy for expression, producing a single cell expression matrix.

Morphological features such as the cell area, eccentricity, as well as the spatial coordinates of each cell can also be measured. PRISM parses these data into an AnnData feature table for further single cell and spatial analysis.

### PRISM Provides a Highly Modular and Interactive Interface for Single Cell and Spatial Analysis

Within the AnnData analysis group, PRISM provides a complete interface for performing single cell and spatial analyses completely using interactive widgets, requiring no expertise in coding (Figure 3). However, manipulating AnnData objects throughout an analysis involves reducing, adding, and conditionally subsetting cells and markers. In the lack of a coding environment, accessing AnnData objects therefore needs an interface. Rather than a mix of objects presented to the user, PRISM sought to organise objects in a particular data structure to accommodate user-driven analysis with only interactive widgets. As an object is analysed, it is modified and reshaped with each computational step. With static pipelines, users therefore face difficulties in exploring different computational and analysis choices. To accommodate this, we introduce AnnDataTrees, which organise AnnData objects in a tree-like hierarchy where nodes act as computational checkpoints, storing the AnnData object at that step. Each object, therefore, points to a parent object representing the object state at the previous step, and child objects represent the subsequent states or computations the user makes on the object. This is especially useful for reducing computational time by removing the need to rerun an analysis pipeline from the very beginning but instead allows iteration from previous checkpoint node. Users interact with this tree in the form of nested rows by clicking on the node that they wish to have loaded into the analysis modules.

**Figure 3:**
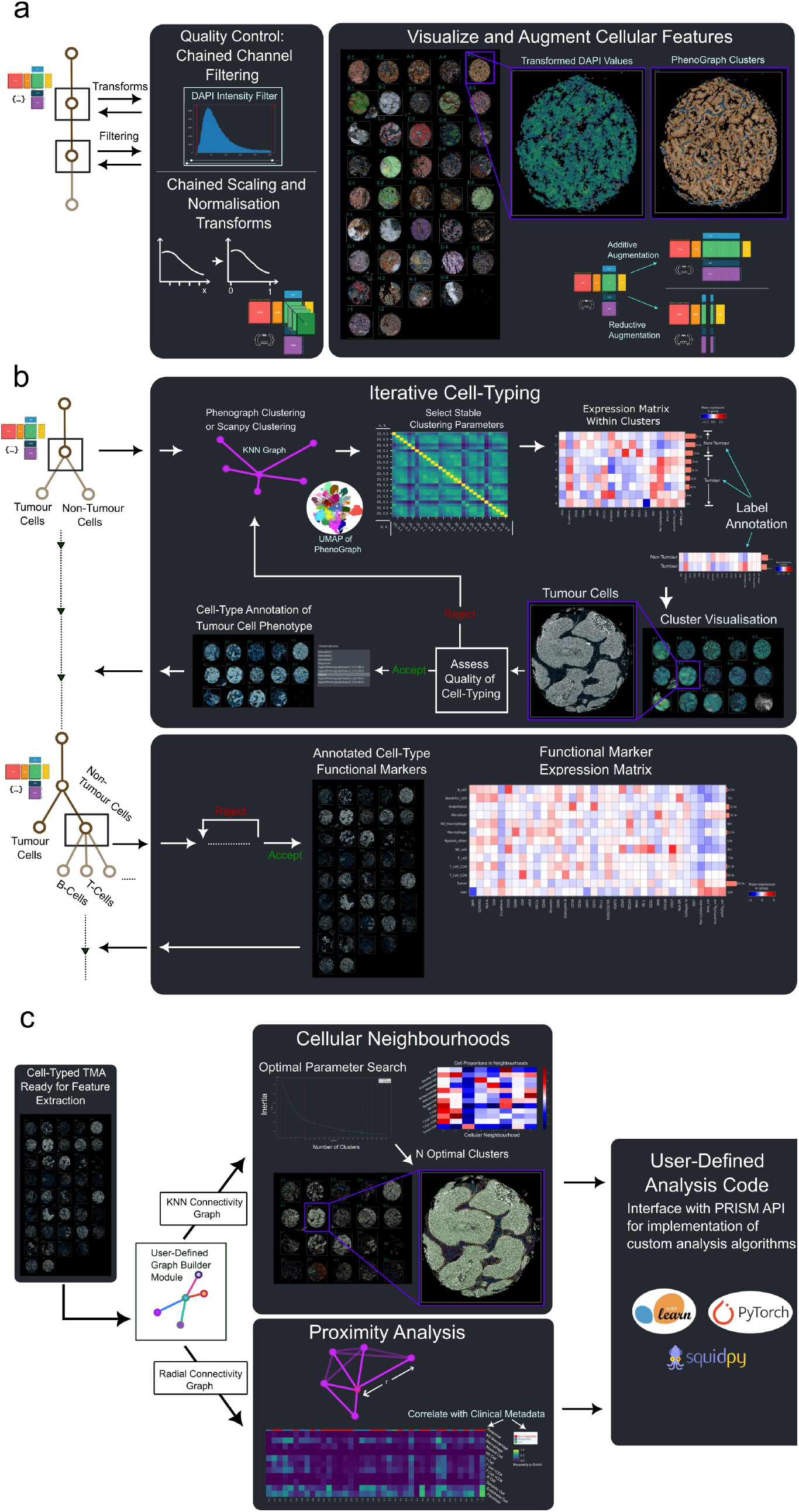
An overview of PRISM’s preprocessing and quality control tools, functionality for iterative cell-typing, and spatial analysis tools. **a)** An overview of the filters and scaling/normalisation that can be chained within the analysis tab for quality control (left panel), and an example of the visualisation of a preprocessed and normalised TMA suitable for downstream cell-typing (right panel). Further data augmentation is possible for the combination and reduction of the feature space (right panel). **b)** An example of the iterative cell-typing process showing sequential tumour/non-tumour cell identification followed by additional cell-typing using marker positivity. PhenoGraph clustering using stable parameter selection, followed by manual annotation of cluster labels and assessment of cell-typing quality, is applied sequentially to assign cell-type labels. **c)** An overview of the spatial analysis tools incorporated into PRISM. The CN assignment is driven by an elbow parameter search to assign classifications to an optimal number of neighbourhoods (middle top panel). Spatial analysis tools such as proximity analysis via p-scores (ref) (middle lower panel) and nearest-neighbour calculations allow for the quick evaluation of cellular patterns within tissue subregions. User-defined analyses can be implemented by use of the API, which wraps a variety of spatial- and data-analysis python libraries (right panel).

The AnnData analysis modules are presented to the user in a sequence of arranged tabs, although the user can typically perform any step, from any module, in any order. These modules involve augmenting (adding or removing) expression features, performing preprocessing, unsupervised clustering with an integrated parameter search and assessment, interactive cell type annotation, and subclustering (Figure 3). The augmentation module allows users to add and/or remove features to the expression space, useful for dropping low quality markers, using specific marker sets for clustering specific cell populations, or adding other features such as morphological measurements to aid clustering. The preprocessing module allow users to perform transformations to normalise and scale data with single chained transforms, performing interactive QC, and filtering based on various metrics within an interactive histogram plot (Figure 3a, left panel). Uniquely, PRISM integrates directly with SpatialData’s dedicated napari plugin, napari-spatialdata, where the user can use the view widget to assess QC and explore every single cellular feature value by visualising them as colormap intensities on the linked cell segmentation mask (Figure 3a, right panel).

Cell-type annotation by unsupervised clustering is a highly iterative process that requires manual assessment of input parameters including chosen markers, the number of neighbours in a high-dimensional embedding space (K), and the resolution value (R) for graph clustering. In the unsupervised clustering modules (Figure 3b), PRISM implements a grid searcher which optimises the process of searching through user provided ranges of K and R for the PhenoGraph [28] and Scanpy [18] based clustering methods. The clustering runs can then be compared and visualised in the parameter assessment module, using metrics such as the adjusted rand index (ARI), adjusted mutual information (AMI) and normalised mutual information (NMI) to determine parameter settings which have relatively stable clustering results. The user can then use this to guide their choice of clustering run(s) from which to extract labels. In the cell type annotation module, the mean expression per cluster label can then be visualised from a selection of heatmap-like plots. Together with the plots, cell types can then be easily annotated using a pop-up annotation table. Cell type annotations can then be further checked by the user on the linked cell segmentation mask (Figure 3b, lower row). Depending on the annotations, the user may choose to cluster differently or further functionalise cell types by using the subclustering module. Figure 3b highlights this iterative process as a fully integrated workflow within PRISM in real-time without reliance on external display tools. To keep the experience as interactive and dynamic for the user, PRISM reduces computational time by implementing GPU accelerated implementations for certain functions and algorithms.

Spatial analyses are typically performed in standalone coding environments after export of cell-typed data to an AnnData object, which limits data visualisation and fast modification of analysis parameters and strategies. In PRISM, neighbourhood identification and spatial proximity analyses can be performed in a fully integrated component. The graph builder module allows the user to construct different types of spatial graphs, such as k-nearest neighbour (KNN) and radial connectivity graphs, which in turn, streamline the downstream analysis process (Figure 3c). Cellular neighbourhoods (CNs) [25] can be defined by a K-Means clustering method applied to a set of neighbourhood histograms of the cell-type KNN graph, and connectivity graphs can be applied to analyse spatial proximities and other similar metrics using annotated cell features (Figure 3c, middle panel). Figure 3c highlights how the cellular composition of these CNs can be easily quantified and visualised. Subsequent spatial analysis of the CN-agglomerated data can then be performed to quantify high-level features such as same-label and multi-label proximity metrics within and between neighbourhoods.

## Discussion

Spatial proteomics has emerged as a powerful approach for characterising the TME and tissue contexture, and together with sampling many tissues through TMAs, enable deep insights into the cellular architectures and mechanisms underlying diseases across large patient cohorts. While multiplexed imaging technologies like CycIF and CODEX/Phenocycler Fusion offer detail on tissue and protein structure, they generate complex and high-dimensional datasets that pose significant computational challenges. The interpretation of these rich datasets is further hindered by the lack of user-friendly and highly interactive analysis tools.

We present PRISM as solution designed to make analysis more accessible and interactive. PRISM empowers users by providing control across all aspects of analysis, from processing raw images to performing spatial analysis with user-generated annotations - all within a single graphical user interface. Recognising that PRISM targets translational and clinical research communities, which include a diverse variety of wet and dry lab researchers, we prioritised making the tool interactive and accessible by implementing every functionality through a graphical interface that eliminates the need for any coding expertise.

To achieve this goal, we innovated in design and integration. PRISM is built on foundations of the SpatialData framework, allowing users to access and store a wide range of data modalities produced throughout analyses. This framework also supports modular computational workflows and is compatible with existing GUI frameworks such as napari. The integration of SpatialData also enables the storage of various data modalities in scalable formats like Dask arrays and Zarr stores. Furthermore, this integration goes beyond mere software compatibility, as PRISM provides an easy way to use such tools with dedicated widgets within the napari viewer. Together with the napari-spatialdata plugin, PRISM harmonises visualisation and analysis of images, regions, shapes and cells in one viewer.

PRISM includes flexible image processing modules that extract clinically relevant structures such as tissue cores and compartments with the masking and de-arraying module. With a highly customisable masking module, users have more control over how image masks are generated, ranging from the channel(s) used, to the resolution they are computed at. This allows users to tailor masks to their use case, whether it be to enhance de-arraying by avoiding noisy or un-important regions of the image or to perform compartmental-based analyses.

PRISM streamlines the process of cell segmentation by providing user-friendly interfaces that allow users to control their desired segmentation strategies and models. A novel feature enables users to merge multiple channels for more robust cytoplasmic-based cell segmentation, as no single cytoplasmic marker ubiquitously stains all cells across the TMA. PRISM adopts Cellpose for cell segmentation due to its various repertoire of models, accommodating both nuclear and cytoplasmic-based segmentation, as well as newer models which address image noise [29]. Additionally, a previewer allows users to verify the quality and validity of the input images prior to optimising cell segmentation, validating if the chosen channel(s) provide a good representation for nuclei, cytoplasm or both.

For analysis of AnnData structures, the lack of a coding environment necessitated a widget whereby users could select a working table to perform analyses on. We therefore introduce the AnnData tree structure to track AnnData objects as computational checkpoints that have pointers to a parent object and/or child objects(s). This is shown intuitively to the user as a widget of nested rows. While this enables greater user-driven exploration of the data, its key limitation is that the structure wraps AnnData objects, and as a result, a copy of every contained AnnData object is stored on disk. However, the cost of the required disk space should not be a limiting factor in the context of the significantly enhanced processing and visualisation times in analysis.

PRISM’s heavily user-guided approach is justified by the complex and varied nature of translational and clinical research, particularly in the context of TMA analysis. TMAs are derived from specific organs and processed using multiplexed immunofluorescence protocols with carefully selected antibody panels. These panels are tailored for research of various disease mechanisms across a range of biological contexts [30]. Newer antibody panels for high-plex proteomic studies are now over 100-plex [7], demonstrating the need for user-guided functions in analysis frameworks to handle the many combinations and edge-cases in research that may appear. For example, PD-1 may be included in cancer immunotherapy studies to understand sensitivity and resistance to anti-PD-1/PD-L1 inhibitor therapies, but not necessarily in projects that want to study antibody drug conjugates. Together, due to the variability in sample composition, marker panels of TMA tissues, and mechanism of description (e.g. the introduction of interactomics data and combinations with other multi-omics imaging data), software such as PRISM that can provide a highly flexible framework for analyses are beneficial to the field of spatial biology.

In summary, we present PRISM as a fast, convenient, and modular framework for end-to-end analyses of multiplexed proteomic data from TMAs. The interoperable nature of the integrated methods and substantial visualisation capabilities of the framework allow for a user-friendly experience in the analysis of multiplexed proteomics data. The framework is equipped with capabilities that extend from high-quality image denoising, TMA masking/gridding, cell segmentation, and feature extraction modules, to AnnData analysis modules that allow for feature QC, data augmentation, unsupervised cell phenotyping, and graph building for spatial analyses. The modular nature of the framework allows for implementation of both existing tools and the integration of developing tools that are available for all steps of the analysis process. PRISM offers a unique set of end-to-end tools that encompass the range of processing steps in a typical analysis of TMA proteomic data, and it is well-situated for the future in the spatial biology field where flexible, interoperable, and user-driven analyses of spatial data will be key to deriving clinical and biological insights.

## Methods

### Datasets

We primarily provide visual examples and processing benchmarks from a previously analysed [12] TMA of 42 cores derived from the resected tissues of a Non-Small Cell Lung Carcinoma (NSCLC) patient cohort. A single core was dropped for demonstration. Multiplexed proteomic imaging was performed on the TMA using the Phenocycler Fusion (Akoya Biosciences, USA). A total of 33 markers were used for demonstration, constituting a typical range of protein markers used in immuno-oncology studies. The TMA consists of 40 cores, with approximately 250k total cells across the cores. This study has Queensland University of Technology (QUT) Human Research Ethics Committee approval (UHREC #2000000494) and University of Queensland ratification.

### Data and Computational Backends

PRISM parses all data as ‘elements’ into a SpatialData object. Raw multiplexed images are stored as multiscale ‘Image’ elements, which are the images in pyramidal image formats, where the same image is stored at decreasing resolutions or scales. All images are parsed as a chunked Dask arrays, and scales are grouped together in the form of xarray [31] DataTrees (and on-disk as Zarr groups). The binary TMA masks and cell segmentation masks are stored in the SpatialData object as ‘Labels’ elements, both as single-scaled chunked Dask arrays. Shapes such as the polygonal representation of the binary TMA masks, circular TMA cores, or rectangular TMA envelopes are stored in the SpatialData object as ‘Shapes’ elements, all as GeoDataFrames, with a column storing the actual shapes, and an additional column storing the TMA grid label for each shape. Tabular data such as the TMA grid labels (representing again to map shapes to instances), and single cell feature expression tables are stored as ‘Tables’ elements, all as AnnData objects. For computations, certain modules and functions are GPU-accelerated (Supplementary). These utilise functions and algorithms from both the base NVIDIA RAPIDS library [32] and the RAPIDS implementation for Scanpy, rapids-singlecell [19].

### Graphical User Interface Frameworks and Widgets

All widgets are created using both the PyQt and magicgui libraries. PRISM is implemented as a napari plugin, making these widgets embeddable beside the main viewer. Where possible, the widgets redirect the computations to launch in a new thread to avoid blocking the event loop in the main GUI thread, allowing the user to continue visualising and interacting with the viewer while computations are running.

### Software Implementation

PRISM is currently implemented as a napari plugin, designed to be run completely within the napari application, alongside the napari-spatialdata plugin for visualisation capabilities.

Future versions of PRISM will include an API to use PRISM functions as well as access data loaded in the napari viewer within coding environments such as Jupyter notebooks. PRISM loosely follows a Model-ViewModel software architectural pattern. The models are the computational modules of PRISM which operate directly on the SpatialData objects. The accompanying widgets for each module act as ViewModels, handling the communication between the underlying SpatialData object, the module and user inputs. Certain widgets communicate with napari-spatialdata widgets to signal that a computation was performed and refresh these widgets to show any new attributes, labels, or metadata.

### Masking

For generating masks, the user has control over 1) the input image and 2) some of the image processing functions. For the input image, the user can choose the resolution to compute the mask, the channel(s) to perform masking, and an optional bounding box created by the user in the GUI to constrict the masking operation spatially. If the user chooses multiple channels, the functions are applied separately to each channel, and then the binary masks are merged. The scikit-image package [33] was used for all image processing functions. The masking function consists of image processing functions that are applied to the raw image in an ordered sequence:

1. (Optional) Li Thresholding [34] on the image to reduce background noise.
2. (Optional) A 2D Sobel kernel [35] to reduce the effects of out-of-focus regions on the masking process by highlighting distinctive cellular boundaries.
3. (Optional) A gamma correction (defaults to gamma = 0.2) for scaling intensity values into a more compressed dynamic range.
4. A three-class Otsu threshold [36] to separate the image into 3 intensity classes, where the image is binarised via thresholding and keeping only the last class.
5. Expansion of the binary masks by a user set number of pixels (microns). Further details on each step can be found in the Supplementary Materials.

### TMA Dearraying

We provide a simplified de-arraying algorithm consisting of 1) the detection of circular-shaped TMA cores or blobs, followed by 2) the estimation and fitting of a TMA grid using those detected blobs. These masks can either be binary masks (for example generated in the masking module) or segmentation masks (of TMA cores with any external method). If segmentation (instance-labelled) masks are provided, the inputs are binarised (all non-zero values converted to 1). The cores are labelled by row using letters and by column using numbers (i.e. the top left most core will be labelled A-1, the core to the right of it will be labelled A-2, and the core below it will be labelled B-1). Further details on these blob detection and grid estimation algorithms are found in the Supplementary Material.

### Cell Segmentation

We provide an interface which allows the user to choose from all available Cellpose models for cell segmentation, including the built-in models (“nuclei”, cyto3”, “cyto2”, etc.), the newer image restoration models added in Cellpose 3.0 (“denoise_”, “deblur_”, “upsample_”, etc.) and custom user-trained models (the interface includes a file dialog widget where users can navigate and open their model folder). Users can choose to enforce tiling by TMA cores or envelopes instead of tiling and segmenting the entire image, and users can provide singular or multiple channels for segmentation. If multiple channels are chosen, they are merged. The user can choose to merge them with a maximum (recommended), median, mean, or summative intensity projection. We provide this functionality as a single cytoplasmic protein may not be expressed across different images, tissues, or cells due to biological or technical issues. Some models such as “cyto3” may also benefit from an additional singular nuclear channel chosen by the user. A nuclear diameter can be provided to assist the model in its predictions. This value defaults to -1, which directs Cellpose to perform an automatic estimation of diameter. The two main tweakable parameters by the user to optimise segmentation results are the “Cell Probability Threshold” and “Flow Threshold”. The user can decrease the former if the model is under-segmenting in dim areas, and/or decrease the latter if the model is segmenting unexpected shapes (will tend to under-segment as it will segment more uniform morphologies). We provide a segment the “First and Last Tile only” option, if tiling by TMA annotation, for quick validation of the segmentation. We also provide an input image previewer to check if the selected channel(s) for segmentation provide a good visual representation of what the user is trying to segment. Segmentation will run on the CPU if the CPU version of PyTorch is installed. GPU-accelerated segmentation will be enabled depending on user hardware. If the user is running OSX (MacOS), the segmentation module will utilise the MPS (metal performance shaders) backend for GPU-accelerated segmentation. If the user has access to a CUDA-compatible GPU and has installed and configured PyTorch with an appropriate CUDA library, the module will run GPU accelerated segmentation by default.

### Expression Measurement

In this module, we provide a way to measure image properties of each cell from the cell segmentation masks generated by the segmentation module. We utilise the “regionprops_table” function from scikit-image for these measurements. With these properties, single cell level information is captured and produces the expression-like tabular data of a single cell experiment. For each cell, the user chooses either the mean or median intensity of each channel or protein as a proxy for expression. Then, additional properties are measured related to the cell’s morphology, such as area, eccentricity, and solidity, as well as each cell’s spatial coordinates. If the user provides a tiling layer, such as the TMA core, this is added as a metadata column to track the originating core of each cell. The output is an AnnData table, with cells as .obs row entries, intensity measurements as .var columns, and metadata in the .obs columns. The spatial coordinates of each cell are parsed to .obsm[“spatial”]. This table is stored in the SpatialData object and is linked to the segmentation mask for downstream cross-referencing.

### AnnData Trees

We introduce AnnData trees, which hold and save AnnData objects in a tree data structure. We did this to track highly interactive and exploratory analyses, where the user can perform many different combinations of operations on many different AnnData subsets. The tree is an instance of the QTreeWidget class from PyQt. The nodes populating this tree are custom classes we develop called “AnnDataNodeQt” that extend the QTreeWidgetItem class from PyQt. AnnDataNodeQt objects contain AnnData objects, as well as a reference to the on-disk path of the object Zarr stores. AnnDataNodeQt instances load parent-child relationships via a dictionary in .uns[“tree_attrs”] that holds references to the on-disk path of its parent and children nodes. Each AnnData object is stored on-disk and on-memory as separate table elements in the containing SpatialData object.

### Augmentation

The augmentation module contains the functions to modify the shape of the expression matrix of the single cell AnnData object. Here, the user can choose to expand the expression matrix feature space by adding numerical columns from .obs as .var attributes. For example, the user can choose to add morphological features like area, eccentricity and solidity as ‘expression’ features. The user can also choose to reduce the feature space by removing features (i.e. due to poor quality markers or performing clustering on a subset of markers). All modifications with this module create an AnnData subset in the AnnData tree.

### Transforms

We provide transform functions in the preprocessing module that can be applied to the expression matrices. The user can chain any number of transforms in any order and combination, tailored to their dataset. We provide 5 transforms:

- Arcsinh transform with a co-factor of 150 applied elementwise.
- Scaling to unit variance and zero mean (from scanpy).
- Percentile transform to the 95^th^ percentile.
- Z-score transform along rows.
- Log transform with a pseudocount of 1 applied elementwise (from scanpy).

### Embeddings

In this module, we provide wrappers for dimensionality reduction and embedding-related Scanpy (and rapids-singlecell) functions. The inputs may either be the expression matrix, or on a reduced embedding matrix. We provide the following functions with all the parameters in the interface:

- Principal components analysis
- t-stochastic neighbours embedding
- K-nearest neighbours (KNN) embedding
- Uniform manifold approximation and projection
- Batch-effect correction and integration of PCA embeddings with Harmony [37]

### Unsupervised Clustering with Parameter Searches

Clustering methods require rather arbitrarily set parameters, which are tweaked by the user until ‘reasonable’ clusters are generated. This can lead to many repetitions of:

1. Clustering
2. Plotting and assessing the clustering results
3. Repeat steps 1-2 with different parameters if results are of poor quality
4. Annotating the clusters based on marker expression and the raw image
5. Repeating steps 1-4 if subclustering these annotated clusters

In this module, we streamline this process by performing steps 1-3 over a range of parameters with custom implementations of clustering methods, thereby putting the plots, raw image and annotation table in the single GUI window. The AnnData tree structure keeps track of the resulting subsets and subclusters.

We have implementations for two similar clustering methods: PhenoGraph and Scanpy. Both methods involve the construction of graphs from an embedding representation of the cell feature matrix, followed by Leiden [38] graph clustering (or community detection). For PhenoGraph, we provide a CPU-only and hybrid implementation utilising a mix of CPU and GPU computations. For the GPU-accelerable parts, we use RAPIDS cuml for constructing the KNN graph on the expression space, and for Leiden clustering on the Jaccard graph. We perform the conversion of the KNN graph to a Jaccard graph on the CPU due to the high memory demand of this computation. For Scanpy clustering, we wrap the Scanpy functions for CPU-only computations and rapids-singlecell functions for GPU-only computations. We implement a parameter searcher for both methods, where many clustering runs over many Ks and resolutions are performed in batch. To optimise a searching across many Ks, we compute the KNN graph once with the largest K and query this graph for smaller Ks. We enforce KNN graphs to be computed using a brute-force algorithm to ensure exact results for querying. We provide to the user a visual indication of reasonable values for K and resolution by assessing their ‘clustering stability’ using a heatmap of clustering quality metrics between a clustering run and all other clustering runs. Currently we have the following metrics for the user’s assessment: adjusted mutual information (AMI), normalised mutual information (NMI), or ARI (adjusted rand index). We compute these metrics by taking the results of one clustering run as the “true labels”, and the other clustering runs as “predicted labels”.

### Spatial Graph Construction

The basis for spatial analyses is defining how cells relate to one another in physical space. These relationships are represented in the form of a spatial graph, where nodes are cells in space and edges are how cells are connected. Different spatial analyses operate with different graphs, and so we wrap the “spatial_neighbors” function from squidpy as 1) a way for the user to construct many different graph types and 2) to standardise spatial graph construction. As PRISM is designed for imaging-based data, we enforce the “coord_type” parameter to be “generic”. The following graphs can be constructed:

- KNN (suitable for characterising neighbourhoods)
- Radial-connectivity neighbours (what distance does the user constitute to be an ‘interaction’ between cells)
- Delaunay (graphs with edges that do not interfere too much, i.e. does this model cover non-blocking / direct interactions, irrespective of distance, and hence cover auto + paracrine signalling)

### Cellular Neighbourhoods

We provide a wrapper for computing cellular neighbourhoods (CNs) [25], directly compatible with the spatial graphs constructed with squidpy. CNs are computed by using the KNN graph to extract the frequency of a cell neighbouring a particular phenotype. A cell by phenotype matrix is used for KMeans clustering, with K being equivalent to the number of CNs identified. Instead of the user providing a single value for K, we provide a parameter search to compute a range of Ks or CNs as previously done [11], and utilise the Kneedle [39] elbow detection algorithm to identify an ‘optimal’ K. We provide the inertia or sum of squared distances of each cell to its assigned CN centroid as a quality metric and compare this with the K value for each run. The ‘optimal’ K can be defined as the lowest possible K with the lowest value for inertia. We identify this ‘elbow’ automatically with the Kneedle algorithm (implemented in python with the “kneed” package [40]), setting the function parameters with the curve set to “convex” and direction set to “decreasing”. The user can visually verify, in a line plot with the elbow point annotated, that the detected ‘elbow’ for K is a reasonable estimate of K against inertia.

### Proximity Density

We provide a wrapper for computing the proximity density component of the “spatial_pscore” function from scimap [21], directly compatible with the spatial graphs constructed with squidpy. Proximity density is defined as the number of instances a given cell phenotype is next to another cell phenotype (as defined by the spatial graph), divided by the union of cells of those phenotypes. We provide an interface for the user to easily define which combinations of cell types to perform the pairwise comparisons. The user can visualise these scores in the form of a PyComplexHeatmap [41], with rows as pairwise comparisons and columns as a user-specific grouping variable, such as each unique TMA core. Each grouping variable can be annotated with an accompanying metadata variable (i.e. annotating each TMA core with patient metadata, such as survival state or patient response to a treatment).

## Code Availability

PRISM is available on the following repositories:

- GitHub: https://github.com/clinicalomx/napari-prism (latest development builds)
- napari-hub: https://www.napari-hub.org/plugins/napari-prism (latest version)

Documentation and tutorials for using PRISM can be found at the following url: https://napari-prism.readthedocs.io/en/latest/

## Data Availability

Data is available upon reasonable request to the corresponding author.

## Acknowledgements

The authors would like to acknowledge that this study was supported by the Passe and Williams Foundation, Cure Cancer, and the Wesley Research Institute.

## Author Contributions

Study design: RT, CWT, JM, AK2

Experimentation: RT, AK1, CWT, JM

Analysis: RT

Development: RT

Writing and critical review: all authors

^*^Aaron Kilgallon (AK1), Arutha Kulasinghe (AK2)

