## Supplementary Material for "PRISM: A Python Package for Interactive and Integrated Analysis of Multiplexed Tissue Microarrays"

**Li Thresholding**

Li thresholding aims to find the optimal threshold, $t$, by minimising the cross-entropy between the original image, $I\left( x, y \right)$ and its binary segmentation. PRISM uses the implementation from scikit-image (“skimage.filters.threshold_li”).

The cross-entropy function $E\left( t \right)$ is defined as:

$$E\left( t \right)= -\sum_{x, y} [p\left( x,y \right)\log q\left( x,y \right)+\left( 1-p\left( x,y \right) \right)\log(1-q(x,y))]$$

Where $p\left( x,y \right)$ is the normalised pixel intensity of the image, defined:

$$p\left( x,y \right)=\frac{I(x,y)}{\sum_{x,y} I(x,y)}$$

And $q\left( x,y \right)$ is the thresholded binarised image, defined:

$$q\left( x,y \right)=\left\{ \begin{aligned} 1, if I\left( x,y \right)\geq t \\ 0, if I\left( x,y \right)<t \end{aligned} \right.$$

Threshold $t$ is iteratively updated with gradient descent:

$$t_{k+1}=t_{k}-\alpha\frac{\partial E}{\partial t}$$

Where $k$ is the iteration, $\alpha$ is the learning rate, and $\frac{\partial E}{\partial t}$ is the gradient of the cross-entropy function with respect to the threshold $t$:

$$\frac{\partial E}{\partial t}=\sum_{x,y} [\delta(I\left( x,y \right)-t)(\log\left( q\left( x,y \right) \right)-log(1-q(x,y)))]$$

With $\delta$(x) representing the Dirac delta function.

Iteration continues until a tolerance, $\epsilon$ is reached:

$$\left| t_{k+1}-t_{k} \right|<\epsilon$$

PRISM uses the default heuristic from scikit-image for setting the tolerance value $\epsilon$, which is set as half the smallest difference between intensity values in $I\left( x,y \right)$).

The image is then thresholded (but not binarised) using 90% of the identified threshold, $t_{optimal}$:

$$I_{thresholded}\left( x, y \right)=\left\{ \begin{aligned} I\left( x,y \right), if I\left( x,y \right)>{0.9t}_{optimal} \\ 0, if I\left( x,y \right)\leq{0.9t}_{optimal} \end{aligned} \right.$$

**2D Sobel Filter**

A 2D Sobel filter aims to enhance edges in the image, $I\left( x,y \right)$by applying a discrete convolution with horizontal and vertical Sobel kernels $S_{x}(x,y)$ and $S_{y}(x,y)$, to get the strength of edges by replacing pixels with the gradient magnitudes. PRISM uses the implementation from scikit-image (“skimage.filters.sobel”).

$$S_{x}= \left[ \begin{matrix} -1 & 0 & 1 \\ -2 & 0 & 2 \\ -1 & 0 & 1 \end{matrix} \right], S_{y}= S_{x}^{T}$$

$$I_{x}\left( x,y \right)=\sum_{u=-1}^{1} \sum_{v=-1}^{1} I\left( x+u, y+v \right)S_{x}(u+1,v+1)$$

$$I_{y}\left( x,y \right)=\sum_{u=-1}^{1} \sum_{v=-1}^{1} I\left( x+u, y+v \right)S_{y}(u+1,v+1)$$

$$I_{e}\left( x,y \right)=\sqrt{I_{x}\left( x,y \right)^{2}+I_{y}\left( x,y \right)^{2}}$$

**Contrast Limited Adapted Histogram Equalisation (CLAHE)**

CLAHE enhances local details, enhancing dark or light parts of a given image, $I(x, y)$. CLAHE is particularly useful for marker channels with uneven staining throughout the image. PRISM uses the implementation from scikit-image (“skimage.exposure.equalize_adapthist”).

**Gamma Correction**

Gamma correction scales the intensity levels, $L$ in an image, $I\left( x,y \right)$ to a compressed dynamic range. PRISM uses the implementation from scikit-image (“skimage.exposure.adjust_gamma").

$$I_{g}\left( x,y \right)=\left( \frac{I\left( x,y \right)}{L} \right)^{\gamma}\times L$$

**Algorithm for Blob Detection**

The detection of circular objects is achieved using the Difference of Gaussians (DoG) on user-provided binary masks, $B(x,y)$. PRISM uses the implementation from scikit-image (“skimage.feature.blob_dog”). In the interface, the user provides a parameter for the expected diameter for each TMA core. This is converted into a radius in pixels, denoted $r$. We then set a minimum and maximum bound for $r$ to determine a set of scales to compute the scale space representation:

$$r_{min}=0.8r$$

$$r_{max}=1.2r$$

We convert these into an approximate equivalent for $\sigma$, where we approximate a 1 pixel radius blob to be $\sqrt{2}\sigma$, therefore, for pixel radius $r$:

$$\sigma\approx\frac{r}{\sqrt{2}}$$

$$\sigma_{min}=\frac{0.8r}{\sqrt{2}}$$

$$\sigma_{max}=\frac{1.2r}{\sqrt{2}}$$

And generate a set of scales $\sigma_{i}$, with sigma ratio $s=1.6$:

$$k=\left[ \frac{\log(\frac{\sigma_{max}}{\sigma_{min}})}{\log(1.6)} \right]+1$$

$$\sigma_{i}=\sigma_{min}\times{1.6}^{i}, for i \in\{0, 1, \ldots, k\}$$

Then given a gaussian kernel $G_{\sigma}$, compute the DoG at various scales as:

$$DoG_{i}\left( x,y \right)=\frac{1}{1.6-1}\left[ \sum_{u=-k}^{k} \sum_{v=-k}^{k} I\left( x-u, y-v \right)G_{\sigma_{i}}\left( u,v \right)-\sum_{u=-k}^{k} \sum_{v=-k}^{k} I\left( x-u, y-v \right)G_{\sigma_{i+1}}(u,v) \right]$$

Then detect blobs by identifying local maxima:

$$DoG_{i}\left( x,y \right)>DoG_{j}\left( x^{'},y^{'}, \sigma_{j} \right) \forall\left( x^{'}, y^{'}, \sigma_{j} \right)\mathcal{\in N}\left( x,y,\sigma_{i} \right),$$

Where $\mathcal{N}\left( x,y,\sigma_{i} \right)$ is the set of neighboring pixels and scales, and $j$ are adjacent scales to $i$. This gives us a matrix of detected blobs, $\mathcal{B=}\left\{ \left( x_{i}, y_{i}, \sigma_{i} \right) \right| i=1, 2, \ldots, n\}$.

**Algorithm for TMA Grid Estimation**

From the matrix of blobs, we estimate the TMA grid by first determining the expected coordinates of each row and each column. We do estimate coordinates based on the availability of user-set parameters. In the widget, the user can provide the number of expected rows and/or columns in the TMA grid. If none are provided, then a 1-D point clustering heuristic issued to estimate the expected coordinates:

Given the matrix of detected blobs computed above, $\mathcal{B=}\left\{ \left( x_{i}, y_{i}, \sigma_{i} \right) \right| i=1, 2, \ldots, n\}$,

$$\sigma_{mean}=\frac{1}{k}\sum_{j=1}^{k} \sigma_{k}$$

Take $x_{i}$ as a vector of points, and $\sigma_{mean}$as the expected distance between segments and apply Algorithm 1.

Algorithm 1: Grid search for TMA cores.

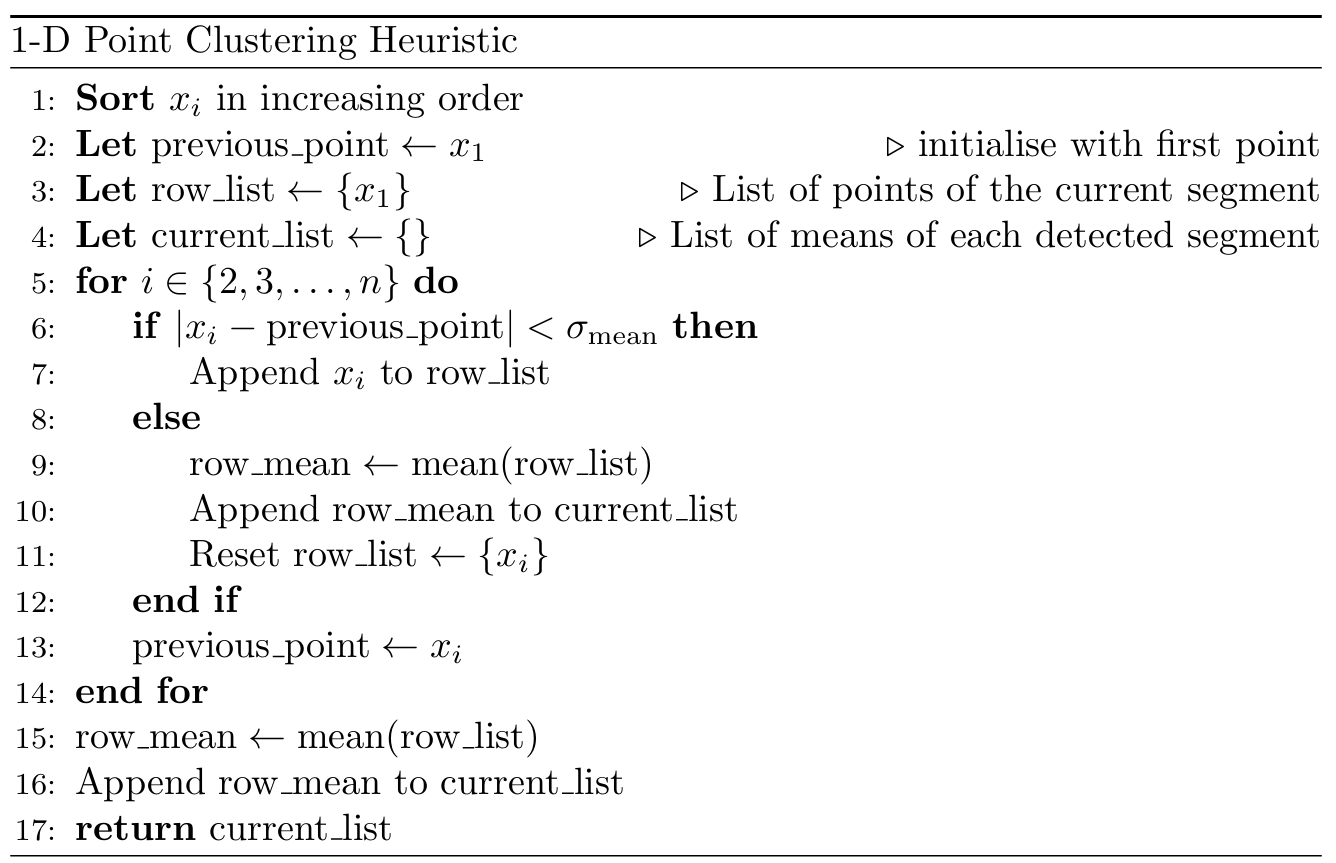

If the user provides the number of expected rows or columns, $k$, then a K-Means clustering algorithm from the scikit-learn package is applied to $x_{i}$, with $k$clusters to generate $current list$, with the values set to the cluster centroids. The above is repeated for $y_{i}$. Given $current list_{x}$ for columns and $current list_{y}$ for rows, we construct a matrix of an ideal TMA grid with expected core positions, $M(x,y)$.
